# Dynamic neutrophil lipidome remodeling during induction of NETosis

**DOI:** 10.1101/2025.10.09.681342

**Authors:** Patrick Münzer, Cristina Coman, Gundula D Lingens, Nina N Troppmair, Jesse A Michael, Reuben S E Young, Stefanie Rubenzucker, Jens Martin, Jann Arden, Melina Fischer, Nils Sülzle, Ferdinand Kollotzek, Monika Zdanyte, Claudia Hornef, Shane R Ellis, Oliver Borst, Robert Ahrends

## Abstract

Neutrophil Extracellular Trap formation (NETosis) affects a wide variety of clinically relevant human diseases. Although lipid remodeling is essential for neutrophil function and membrane rupture during NETosis, the neutrophil lipidome and its dynamics have not been characterized.

Thus, we establish the first quantitative lipidome of human neutrophils comprising 1,039 species across nine orders of magnitude and map its remodeling during NETosis. NET formation caused profound alterations in the phosphatidylinositol, phosphatidic acid, diacylglycerol and lysoglycerophospholipid levels. Calcium- and reactive oxygen species-dependent NETosis pathways displayed distinct lipidomic trajectories, yet converged on the significance of phospholipid lipase networks. Pharmacological inhibition of this networks altered lipid composition and markedly impaired NETosis, while diacylglycerol (DG) treatment revoked the effect. Altogether our findings reveal lipid remodeling as a fundamental determinant of NETosis and identify interconnected and dependent phospholipid lipase networks with downstream DG-dependent signaling as a potential therapeutic target in NET-associated diseases.

## Introduction

Polymorphonuclear neutrophils (PMNs) represent the most abundant subset of immune cells and mediate important antimicrobial activities. Amongst other mechanisms, neutrophils can expel decondensed chromatin structures designated Neutrophil Extracellular Traps (NETs). First described in 2004 as a pivotal part of the innate immunity (*1*), the formation of NETs (NETosis) has since been linked to the development and progression of a wide variety of clinically relevant human disorders. Beyond their involvement in generating autoantibodies in rheumatoid arthritis and heparin-induced thrombocytopenia (*2, 3*), NETs are highly implicated in arterial and venous thrombosis (*4, 5*), myocardial ischemia/ reperfusion injury (*6*), infective endocarditis (*7*), and myocarditis (*8*). Moreover, under pathophysiological conditions NET formation can be triggered by diabetes (*9*) and cancer (*10*), thus contributing to cancer progression and metastasis (*11*). Despite the remarkable clinical significance, the complex cellular mechanisms underlying NETosis remain ill-defined.

NET formation is a strictly regulated sequence of cellular events leading up to the disassembly of nucleosomes by peptidylarginine deiminase 4 (Padi4)-(*12, 13*) and myeloperoxidase (MPO)-(*14*) dependent mechanisms. Ultimately, entropic chromatin swelling culminates in the rupture of the nuclear envelope and the plasma membrane (*15*) with subsequent expulsion of neutrophilic chromatin together with pro-thrombotic and pro-inflammatory factors (*16*). Effective inducers of these processes in vitro are the calcium ionophore ionomycin and phorbol 12-myristate 13-acetate (PMA), which activate Padi4 and trigger the production of reactive oxygen species (ROS) with subsequent MPO activation, respectively (*17*). Considering the rupture of the neutrophilic plasma membrane may occur hours after NET-induction and an observed gradual increase in plasma membrane permeability (*18, 19*), a highly regulated dynamic shift in the lipid composition of the plasma membrane likely contributes significantly to NETosis. However, a detailed description of the quantitative lipidome and dynamic lipid profile in neutrophils especially upon NET induction is still missing.

Besides a study on the importance of cholesterol for neutrophil function (*20*), research on the exact distribution of specific phospholipid classes remains insufficient. Whereas some studies identified phosphatidylcholine (PC) as the most abundant class in neutrophils and phosphatidylserine (PS) and phosphatidylethanolamine (PE) at similar levels (*21*), others did not analyze PS levels at all and report PE as the most prominent lipid class in neutrophils (*22*). Despite these discrepancies, there is a broad consensus that the lipidome of neutrophils is distinct and characterized by an abundance of vinyl ether and ether lipids, believed to be oxidized and halogenated (*23*), thus providing protection against oxidative stress, including ferroptosis. Moreover, it seems neutrophils contain a diverse array of glycosphingolipids, such as gangliosides and members of the (neo)lacto-series, which appear to be absent in other immune cells (*24*), and may contribute to pathogen recognition and interactions with the humoral immune system. Thereby, especially the glycosphingolipid lactosylceramide (LacCer) seems to be highly prevalent in lysosomal granules of neutrophils (*25*), possibly affecting the clearance of pathogens in line with the innate immune response.

To date, most of the studies on neutrophil lipids have merely examined broad categories, often omitting entire classes or capturing only a fraction of the lipidome. Detailed and quantitative analyses of the molecular lipid species comprising the neutrophil lipidome and highlighting the plasma membrane composition are still lacking and none of the recent studies investigated dynamic lipid changes in neutrophils upon induction of NETosis. Thus, this study aimed to characterize for the first time in a quantitative lipidomics approach the lipid profile of neutrophils and the dynamic changes during NETosis. Consequently, we were able to quantitatively define the most prominent lipid classes in neutrophils and illustrate dynamic lipid changes and pertinent lipid-driven signaling pathways essentially mediating the cellular process of NETosis

## Materials and Methods

### Materials and standards

Acetonitrile (ACN), methanol (MeOH), formic acid and water were purchased in LC−MS grade from Biosolve (Valkenswaard, The Netherlands). Ammonium acetate (NH4Ac), ammonium formate (AF), chloroform (CHCl3), phosphoric acid (PA), potassium chloride (KCl), sodium chloride (NaCl), sodium dodecyl sulfate (SDS), dithiotreitol (DTT), trifluoroacetic acid (TFA), iodoacetamide (IAA), triethylammonium bicarbonate (TEAB) and tert-butyl methyl ether (MTBE) were obtained from Sigma-Aldrich (Steinheim, Germany) and acetic acid (HAc) from Carl Roth (Karlsruhe, Germany). Isopropanol (IPA) was purchased from Merck (Darmstadt, Germany), and tris(hydroxymethyl)-aminomethane (Tris) from Roche Diagnostics (Mannheim, Germany). All lipid standards were purchased from Cayman Chemical (Ann Arbor, USA) or Avanti Polar Lipids (Alabaster, USA) and Labeled carnitine standards set B was purchased from Euriso-top (Saarbrücken, Germany).

### Ethical issues

All blood donors gave informed consent. The study was approved by the institutional ethics committee (032/2024BO2) and complies with the declaration of Helsinki and good clinical practice guidelines.

### Neutrophil isolation

Human neutrophils were isolated employing gradient centrifugation as previously described (*26*). Concisely, EDTA-anticoagulated blood was layered on Histopaque-1119 and spun for 21 min. at 1100g without deceleration. After a washing step, the resulting cell pellet was resuspended in HBSS and layered on a 85%/80%/75%/70%/65% Percoll gradient column before being centrifuged for 21 min. at 1100g without deceleration. The resulting neutrophil layers were washed again with HBSS and subsequently resuspended in phenol-red free RPMI medium containing 10mM HEPES. Neutrophil concentration and purity were determined using a hemocytometer (Sysmex KX-21N).

### Preparation of neutrophils for quantitative lipidomics

The desired cell density (2-3 x 10^4^/µL) was adjusted using phenol-red free RPMI medium containing 10 mM HEPES. 2-3 x 10^6^ cells were stimulated in the absence or presence of PLD inhibitor (FIPI, 1 µM) at the indicated time points. After stimulation, samples were spun down for 10 min. at 400 g and the pellet as well as the supernatant were immediately frozen in liquid nitrogen and stored at −80°C until further processing.

### In vitro NET assay

In vitro NET assays were performed as described previously (*26*). Concisely, isolated human neutrophils were adjusted to a cell density of 3 x 10^2^/µL using phenol-red free RPMI medium containing 10 mM HEPES and 0.05% bovine serum albumin (BSA). 3 x 10^5^ cells were added on coverslips to a 24-well plate with the DG analogon (diC_10_) in the absence or presence of PLD inhibitor (FIPI) for 30 min. at 37°C and 5% CO_2_. Afterwards, cells were stimulated for 4 hours at 37°C and 5% CO_2_ with ionomycin (4 µM) or PMA (100 nM) and subsequently fixed with 1.3% paraformaldehyde (PFA) for 30 min. at room temperature. After washing with PBS, permeabilization and blocking, samples were stained overnight with primary antibodies against citrullinated histone H3 (#ab219407, 1:1000) and neutrophil elastase (#GTX72042, 1:1000). Following two washing steps, the corresponding secondary antibodies (#A31574 and #A21428, 1:1500) were incubated for 2 hours at room temperature and after two additional rounds of washing the samples were finally mounted using ProLong^TM^ Diamond Antifade with DAPI (#P36962).

For analysis of NET formation, at least 3 microscopic images were taken from each sample and the percentage of NET formation was calculated using image J.

### Lipid extraction

Neutrophil pellet and supernatant were used for lipid and protein extraction following the SIMPLEX protocol previously described by Coman et al. (*27*). In short, 225 μl of methanol and the internal standard mixture (Mouse Splash, CerMix II, chol-d7, carnitine set B and an in-house mixture containing odd-chain LPI, LPG, LPA, LPS, AA-d11, DHA-d5, EPA-d5, 13-HODE-d4, 12-HETE-d8, 15-HETE-d8, 9-HODE-d4, LTB4-d4, PGD2-d4, PGE2-d4)) were added to the samples and subjected to ultrasonication for 10 s on ice water. Next, 750 μl of MTBE was added, and samples were incubated for 1 h at 950 rpm at 4°C. To induce phase separation, 188 μl of water were added. After a 10 min centrifugation step at 10,000 g at 4°C, the upper organic phase (containing GPs, GLs, SPs and STs) was carefully removed and dried under a gentle nitrogen flow. To complete protein precipitation, 527 μl of methanol were added to the lower aqueous phase, and samples were stored for 2 hours at −20°C, following centrifugation for 30 min. at 12,000 g at 4°C. The lower aqueous phase (containing FAs) was collected and dried under a gentle nitrogen flow. The protein pellet was dried, resolubilized in 1% SDS buffer, 150 mM NaCl and 50 mM Tris, pH = 8.5 and subjected to the BCA assay.

### Lipid measurements and analysis

#### Shotgun lipidomics

Lipid extracts were resuspended in IPA/MeOH/CHCl3 (4:2:1, v/v/v) with 7.5 mM ammonium acetate and then infused via the TriVersa NanoMate ion source (Advion Biosciences) into an Exploris 240 (Thermo Fisher Scientific) mass spectrometer. The following settings were used for positive and negative mode: ionization voltage +1.25 kV/ −1.25 kV, backpressure 0.95 psi, ion transfer capillary temperature 250°C, S-Lens level of 60% and EASY-IC was enabled. In both modes, full MS spectra were acquired with a resolution of 240,000 followed by a data independent acquisition (DIA) for precursor masses at an interval of 1.001 Da. MS/MS spectra were acquired with a resolution of 60,000 and the precursor isolation window was 1 Da.

#### LC-HRMS analysis for GPLs and GLs of PMN release

5 µl lipid extract resuspended in BuOH:IPA:H2O 8:23:69 (v/v/v) + 25 nM CUDA were loaded onto a YMCAccura Triart C18 column (150 × 2.1 mm, 1.9 μm particle size, YMC Europe, Dinslaken, Germany). Mobile phase A was ACN/H2O (60:40, v/v), mobile phase B was IPA/ACN (90/10, v/v), and both contained 10 mM ammonium formate and 0.1% formic acid. The temperatures of the autosampler and the column oven were set to 8°C and 60°C, respectively and separation was carried out with a flow rate of 0.26 ml min−1 with the following 30-min-long gradient: initial, 30% B; 0.0–3.0 min, hold 30% B; 3.0–15.0 min, ramp to 75% B; 15.0–17.0 min, ramp to 100% B; 17.0–30.0 min, to 5% B; 30.1–35.0 min, to 30% B. The injector needle was automatically washed with 30% B.

The LC was coupled to an Exploris 240, and data were acquired both in positive and negative ion mode. The following electrospray ionization (ESI) source parameters were applied: spray voltage, 3.4 or 2.4 kV, respectively; capillary temperature, 275°C; sheath gas flow rate, 45; auxiliary gas flow rate, 20; auxiliary gas heater temperature, 350°C; S-lens RF level, 60. Full MS spectra from 400 to 1,230 m/z and from 400 to 1600 m/z were acquired in positive and negative mode, respectively with a resolution of 120,000, an AGC target of 1e6, and a maximum IT of 105 ms in one run. DDA with a top 10 experiment was used on the pool samples and acquired in separate runs for positive and negative mode. For MS/MS, a resolution of 15,000, an AGC target of 1e5, a maximum IT of 105 ms and a nCE of 24 and 28 for positive and negative mode were applied.

#### Targeted lipidomics for SLs and STs

Each sample was resuspended in BuOH:IPA:H2O 8:23:69 (v/v/v) + 25 nM CUDA, and 7 µl were injected into the LC system. Analysis of SP and ST was performed as previously described by Peng et al. (*28*) and Troppmair et al. (*29*). Inclusion lists for targeted measurements were generated using LipidCreator (v1.2.0). Briefly, a Vanquish Flex UHPLC system (Thermo Fisher Scientific, Germering, Germany) was equipped with an Ascentis Express C18 main column (150 mm × 2.1 mm, 2.7 μm, Supelco) and fitted with a guard cartridge (50 mm × 2.1 mm, 2.7 μm, Supelco) in a column oven with a temperature of 60°C. Eluent A was ACN/H2O (6:4, v/v; 10 mM AF, 0.1% FA, 5 μM PA), and eluent B was IPA/ACN (9:1; v/v; 10 mM AF, 0.1% FA, 5 μM PA). The separation was carried out at a flow rate of 0.5 ml/min with the following 25 min long gradient: initial (30% B), 0.0–2.0 min (hold 30% B), 2.0–3.0 min (30–56.1% B), 3.0–4.0 min (56.1–58.3% B), 4.0–5.5 min (58.3–60.2% B), 5.5–7.0 min (60.2–60.6% B), 7.0–8.5 min (60.6–62.3% B), 8.5–10.0 min (62.3–64.0% B), 10.0–11.5 min (64.0–64.5% B), 11.5–13.0 min (64.5–66.2% B), 13.0–14.5 min (66.2–66.9% B), 14.5–15.0 min (66.9–100.0% B), 15.0–19.0 min (hold 100% B), 19.0 min (5% B), 19.0–22.0 min (hold 5% B), 22.0 min (30% B), 22.0–25.0 min (hold 30% B). The LC system was coupled to a QTRAP 6500+ (Applied Biosystems, Darmstadt, Germany). The measurements were performed in positive mode with the following ESI source settings: curtain gas 30 arbitrary units, temperature 250°C, ion source gas I 40 arbitrary units, ion source gas II 65 arbitrary units, collision gas medium; ion spray voltage +5500 V, declustering potential +100 V, entrance potential +10 V, and exit potential +13 V. For the scheduled SRM, Q1 and Q3 were set to unit resolution. The scheduled SRM detection window was set to 2 min, and the cycle time was set to 0.5 s. Data was acquired with Analyst (version 1.7.2; AB Sciex) and Skyline(*30*) was used to visualize results and manually integrate signals.

#### Targeted lipidomics for FAs, CerP, lyso-GPs and MGs

The analysis was performed as previously described by Rubenzucker et al. (*31*). The samples were resuspended in BuOH:IPA:H2O 8:23:69 (v/v/v) + 25 nM CUDA and 7 µl were injected onto a YMCAccura Triart C18 column (150 × 2.1 mm, 1.9 μm particle size, YMC Europe, Dinslaken, Germany) fitted with a Vanquish MP35N passive preheater. The column compartment and autosampler temperatures were 45°C and 8°C, respectively. Eluent A was H2O/ACN 80/20 (v/v) and eluent B consisted of IPA/ACN/H2O 60/35/5 (v/v), both containing 0.5 mM NH4Ac and 0.2% HAc. The separation was carried out at 0.4 mL/min with the following 20 min gradient: 0−1 min (hold 30% B), 1−11 min (30-100% B), 11-16 min (hold at 100% B), 16.1-20 min (return to 30% B and hold for re-equilibration). The measurements were performed in both positive and negative mode with the following ESI source settings: curtain gas 35 arbitrary units, temperature 350°C, ion source gas I 45 arbitrary units, ion source gas II 60 arbitrary units, collision gas medium; ion spray voltage +5500 V and −4500 V, respectively. For the scheduled SRM, Q1 and Q3 were set to unit resolution. The cycle time was set to 1 s, the settling time was set to 20 ms and the minimum dwell time was set to 10 ms. Data was acquired with Analyst (version 1.7.2; AB Sciex) and Skyline (*30*) was used to visualize results and manually integrate signals.

### Lipid identification and quantification

All spectra were imported by LipidXplorer (1.2.8) into a MasterScan database under the following settings: mass tolerance 5 ppm; m/z range of 400–1050 and 400-1200 for positive and negative mode, respectively; minimum occupation of 0.15; intensity threshold 5 × 10^3^. Lipid identification was carried on as described by Herzog et al. (*32*) by matching the m/z of the monoisotopic peaks to the corresponding elemental composition constrains. The Molecular Fragmentation Query Language (MFQL) queries were compiled to match precursors and fragment ions in order to confidently recognize lipid species. LipidXplorer output was processed by lxPostman (LIFS) and in-house developed R scripts (RStudio 23.09.1+494). All signal intensities were normalized to the corresponding deuterated internal standard. Protein concentrations, determined by the Pierce BCA Protein Assay Kit, were used to normalize all lipid species, plasma samples were normalized to volume. TGs and CLs were quantified based on precursor intensities.

SPs (Cer, HexCer, Hex2Cer, Sulfo-HexCer (SHexCer), LSM, LCBP, LCB and SM), STs (ST and SE), oxylipins, fatty acids, endocannabinoids, MGs and selected lyso-GPs were identified and quantified by LC–MS analysis. Peak integration from targeted measurements was performed using Skyline (v21.1.0.146). Lipid species abundance was calculated using peak areas and quantified according to the respective internal standard (Ceramide/Sphingoid Internal Standard Mixture II; cholesterol-d7) or as described in Rubenzucker et al. (*31*).

The LC-HRMS data analysis of the PMN release was based on the identification list obtained from the shotgun measurements of the PMNs. The lipid species specific fragments were generated by LipidCreator and monitored in Skyline for peak integration. The MS2-based identification was performed on the pool samples of PMNs, PMN release and plasma which were acquired in the DDA mode, and the lipid species were finally quantified based on the MS1 level. The peak areas were normalized to the ones of the corresponding class-specific standards. All lipid quantities were normalized to the cell number in a KNIME (*33*) or R workflow.

### Isomer resolved Mass spectrometric analysis

Dried lipid extracts were shipped on dry ice from Vienna, Austria to Wollongong, Australia in 200 µL glass inserts housed in amber-glass vials. Upon arrival, samples were stored at −80°C until analysis. On the day of measurement, extracts were equilibrated to room temperature and reconstituted in 20 µL of 4:1 isopropanol:methanol (IPA:MeOH). Aliquots were transferred into a 96-well plate (TwinTec, Eppendorf) and mixed with 2.5 mM sodium acetate in methanol at a 1:1 (v/v) ratio to prepare for nanoelectrospray ionisation (nESI). Ionisation was performed using a TriVersa Nanomate (Advion) operated at 1.4 kV spray voltage and 0.4 psi gas pressure. The ion source was coupled to a modified Orbitrap Fusion Tribrid mass spectrometer (Thermo Fisher Scientific) that allowed passive introduction of ozone into both the high-pressure dual ion trap and the ion routing multipole. Ozone was generated using a Titan-30UHC high-concentration ozone generator (Absolute Ozone) via dielectric barrier discharge of high-purity oxygen, yielding inline ozone concentrations of 220 g/Nm³. Data were acquired using a data-independent acquisition (DIA) method adapted from Michael et al. (*34*). Positive ion mode was employed with the following scan events: (i) full MS1; (ii) MS³ collision-induced dissociation/ozone-induced dissociation (CID/OzID) triggered by neutral loss of the phosphatidylcholine headgroup ([M+Na–183.0661]+); (iii) MS³ CID/OzID triggered by neutral loss of the phosphatidylethanolamine headgroup ([M+Na–141.0191]+); and (iv) MS² OzID. CID/OzID events were stepped in 2 Da intervals across m/z 574–1036 with a 1 Da isolation window. At the MS² level, precursor ions were subjected to a normalized collision energy (NCE) of 45 V for 5 ms, and the resulting headgroup-loss product ion was reisolated for MS³ analysis (dual ion trap ozonolysis, NCE = 0 V, 400 ms). MS² OzID scans were stepped through 2 Da intervals across m/z 650–950 with a 1 Da isolation window, followed by ion routing multipole ozonolysis for 2.5 s at NCE = 1 V.

### Data analysis of lipid double bonds and processing thresholding

For sn-isomer species data was analysed as recently described. (*34*). For double bond isomers data was extracted from the Thermo .raw files using Xcalibur™ v 3.0.63 for selected lipid species and then processed manually using Excel. Data is presented as the percentage contribution of double bond-specific fragment intensity compared to fragments from all isomers. Given the highly complex nature of the lipid extracts, isobars were expected to be prevalent within the mass spectrum, so precursor ions were specified based on theoretical m/z values and then characteristic OzID neutral loss fragments were calculated using a 5 ppm threshold for assignment. The aldehyde and Criegee ions needed to reach a combined signal-to-noise ratio of 3:1 to be identified and the assignment needed to be present in 2 of 3 sample replicates to be included in the graphical visualisations. Additionally, standard error of the mean (SEM) with a 95% confidence interval was calculated per assignment and if error exceeded the calculated mean value, the assignment was discounted.

For analysis of sn-isomers, MS3 scans in the .raw files were processed through the ALEX123 pipeline from Michael et al. (*34*). The list of validated sn-isomers is generated algorithmically from all combined treatment and control files considering appropriate CID/OzID fragments detected at sufficient frequency and accurate mass (+/-2.5 ppm following file-dependent alignment). The ‘’canonomer’’ label refers to phospholipid isomer where the longer (or if identical length, more unsaturated) of two acyl chains is in the sn-2 position.

### Visualization and statistical analysis

For all statistical analyses and plotting of the data, we used in-house scripts generated with R v4.3.1 (https://R-project.org). Statistical differences between 2 groups were evaluated by 2-tailed, unpaired t-test. One-way ANOVA was used for comparing multiple groups. For all p-values, a significance threshold of pL=L0.05 was chosen.

## Results

### Characterization of the quantitative neutrophil lipidome

To quantitatively profile the lipidome of naïve circulating primary neutrophils, we employed a multiplatform lipidomics workflow (Fig. 1a). In unstimulated neutrophils, the most abundant lipid categories, in descending order, were phospholipids (GP), sterols (ST), fatty acids (FA), glycerolipids (GL), and sphingolipids (SP) (Fig. 1b). Within GPs, phosphatidylcholine (PC), phosphatidylethanolamine (PE), and phosphatidylserine (PS) predominated, whereas phosphatidylinositol (PI), phosphatidylglycerol (PG), and phosphatidic acid (PA) were comparatively minor (Fig. 1b, c). Interestingly, over 50% of the entire class of PE and PC is covered by ether lipids. Neutrophils exhibited a lipidomic dynamic range spanning nine orders of magnitude (Fig. 1c, e), with cholesterol (32743 pmol/million cells) as the most and prostaglandin E_2_ (PGE_2_) as one of the least (0.0021 pmol/million cells) abundant molecules in PMNs (Fig. 1e), whereas cholesterol (2538 pmol/million cells) and LSM (0.0022 pmol/million cells) present as the highest and amongst the lowest levels within the PMN release (Fig 1e). Compared with megakaryocytes, neutrophils displayed a less complex lipid repertoire, yet were more complex than platelets, where 30 species account for ∼90% of the lipidome (Fig. S1) (*35, 36*). The distribution among classes and species is comparable between the PMN membrane lipidome and secreted lipids (Fig. 1 d-f, S2a), with the exception of a lower lipid amount (∼ 4% of the PMNs lipidome), less species and increased levels of cholesterol and hexosylceramides detected in the PMN release. Importantly, upon lipid release, the lipid composition of vesicular membranes diverged from that of plasma membranes, indicating a selective remodeling of the neutrophil lipidome during vesiculation. Notably, neutrophil complex lipids consisted primarily of the ten most abundant fatty acids (Fig. 1g), including 16:0, 18:0, 18:1, 18:2, and 20:4 and the sn positional and double bond isomer analysis of the most abundant PCs and PEs displays species specific distributions (Fig. S3, 4).

**Figure 1.**
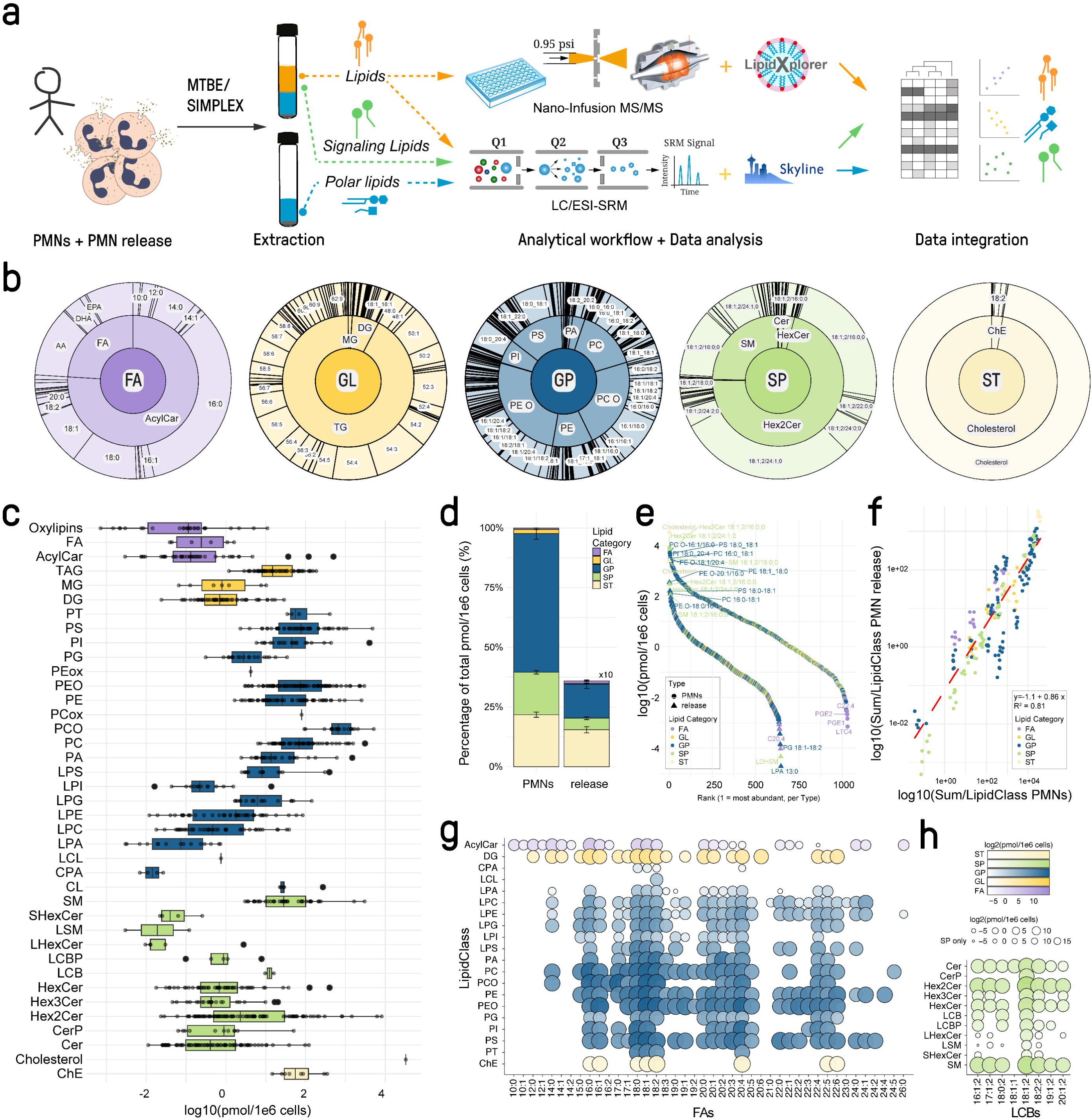
Quantitative lipidomics reveals a complex and multifaceted lipidome for polymorphonuclear neutrophils covering a wide, dynamic range spanning 9 orders of magnitude. (a) Quantitative lipidomics platform covering membrane lipids, signaling lipids and lipids for energy demands. (b) Displays the quantitative distribution of the five lipid categories analyzed in mol% per category; Inner circle illustrates class distribution and the outer circle shows the distribution of species within each class. (c) Boxplots presenting the dynamic range of the PMN lipidome at the resting stage, where each entry represents a lipid species. Quantities of five lipid categories (fatty acids, sphingolipids, glycerolipids, glycerophospholipids, and sterols) spanning 37 lipid classes and 1042 lipid species over a concentration range of nine orders of magnitude are shown. Each bar is composed of all quantified lipid species within the respective class. SM, sphingomyelin; SPBP, sphingoidbases-phosphate; SPB, sphingoidbases; HexCer, hexosylceramide; Hex2Cer, dihexosylceramid; Cer, ceramide; TG, triacylglycerol; DG, diacylglycerol; PS, phosphatidylserine; PI, phosphatidylinositol; PEO, phosphatidylethanolamine-ether; PE, phosphatidylethanolamine; PCO, phosphatidylcholine-ether; PC, phosphatidylcholine; PG, phosphatidylglycerol; PA, phosphatidic acid; LPA, lyso-PA; LPI, lyso-PI; LPG, Lyso-PG; LPE, Lyso-PE; LPC, Lyso-PC; CL, cardiolipin; ST, cholesterol; SE, cholesterol-ester.(d) Comparison of the quantitative composition of the PMN cell and its lipid release at the resting state in mol%; Colors represent categories. (e) S-Plot of the cellular and PMN release lipid concentrations displaying more and less abundant molecules across nine orders of magnitude. (f) Linear regression demonstrating similar patterns at lipid class level between PMNs and its release. (g) Fatty acyl distribution across different classes including GL, GP, ST and Acyl-carnitines. (h) The sphingolipids’ long-chain base distribution for the analyzed sphingolipid classes. All data represent mean values of 6 biological replicates.

Interestingly, levels of bound docosahexaenoic acid (DHA) and eicosapentaenoic acid (EPA) were noticeably lower in neutrophils compared with naïve B cells, megakaryocytes, or platelets (*36*), indicating a limited capacity for producing lipid mediators derived from these precursors (Fig. S2 b-c). Sphingolipid analysis revealed a high diversity of long-chain bases (LCBs), including the major LCBs 18:1, 18:2, 17:1, 16:1 and 18:0 (Fig. 1h) while the bound fatty acids ranged from 14:0;0 to 26:1;0 (Fig. S2 d-e).

Together, these results establish a comprehensive and detailed reference lipidome for circulating naïve neutrophils. They illustrate an unprecedented lipid architecture, characterized by ether lipid enrichment, limited DHA/EPA availability and distinct remodeling upon lipid release – features that may significantly shape the distinct inflammatory and signaling function of neutrophils. The novelty and magnitude of these findings will surely pique the interest of the scientific community.

### Characterization of the lipidome upon NET induction with ionomycin

To investigate the dynamic lipidome in neutrophils upon NET induction, primarily isolated neutrophils were stimulated for 1, 30, and 120 min. with well-established NET inducers ionomycin or phorbol 12-myristate 13-acetate (PMA).

During ionomycin treatment (Fig. 2a), the calcium ionophore increases intracellular Ca²L levels via transportation across plasma and organelle membranes. In neutrophils, this leads to the activation of the calcium-dependent effector Padi4 with subsequent citrullination of histones and finally DNA decondensation. In general, increasing intracellular Ca^2+^ levels affect downstream calcineurin–NFAT signaling and, most likely, phospholipase A_2_ (PLA_2_) activation. However, apart from fatty acyls and lyso-phospholipids, most lipid categories in PMNs are not displaying noticeable shifts within 2 hours of activation (Fig. 2b). The most abundant classes, including PC and PE, as well as less abundant classes, such as PS and PG, are not regulated within the first 30 minutes (Fig. 2d). The exception lies in the downregulation of PI by 50 %, leading to increased levels of LPI both inside and outside the cell, likely through PLA_2_ activity (*37*). This activity is underscored by higher intra- and extracellular levels of LPC, LPS, LPE, LPG, and LPA (Fig. 2h), whereas lyso-sphigomyelin levels remain unchanged in PMNs (Fig. 2d, right). PLA_2_ activity drives an overall increased production of free AA, DHA, DPA and EPA (Fig. 2 e-f and Table S1) as the most abundant PUFAs in neutrophils, correlating with a rapid upregulation of oxylipins within the first minute (Fig. 2g) (*38*). The comparatively minute impact of PLA_2_ on highly abundant PUFA is likely related to the presence of ethers instead of esters in 2/3 of all PLs, reducing the accessibility towards PLA_2_. Notably, all PLA_2_-dependent changes occur within the first minute of activation (Fig. 2 f-h). However, as a large contributor to AA release the precursor PI 18:0_20:4 was identified similarly as in platelet activation (Fig. 2n) (*36*).

**Figure 2.**
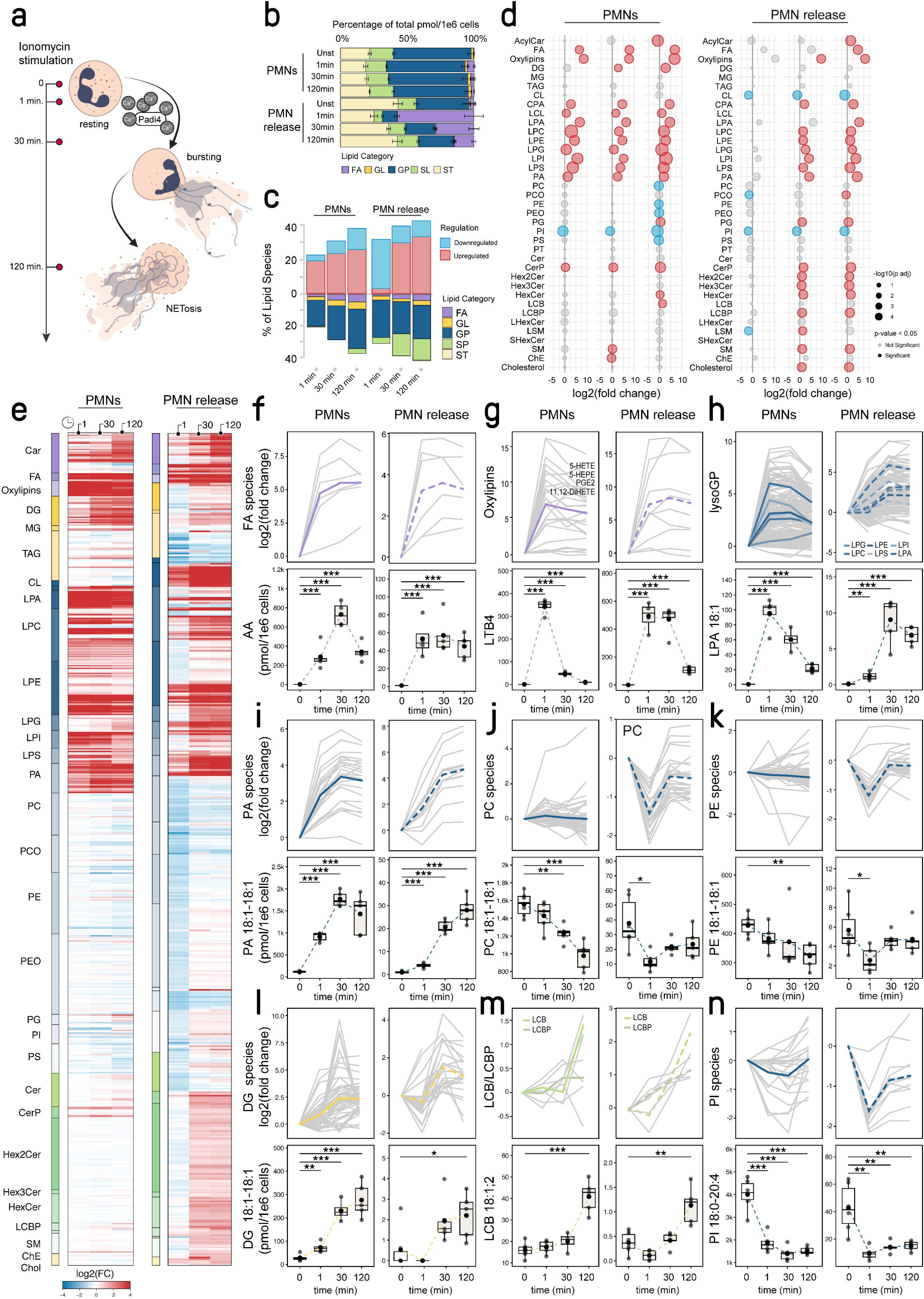
A rapid dynamic lipidome remodeling of PMNs is triggered by Ca^2+^ influx during NET-formation. (a) Timeline of sampling after NET induction using 4 µM ionomycin triggering Ca^2+^ influx. (b) Category-based lipidome remodeling of PMNs and release after NET-induction showing glycerophopholipids, glycerolipids sphingolipids, sterols and fatty acyls. (c) Quantitative assessment of the number and direction of species regulated in each class during NETosis at 1, 30, and 120 minutes. (d) Ratio plots presenting class dynamics at 1, 20 and 120 minutes (left to right) for the cellular and the released lipidome. The ratios are depicted as log2 fold changes. Here, the most prominent modulations of the lipidome at class level during NET-formation and release are displayed. Significant changes are colored in red for up- and blue for downregulated classes. (e) Heatmap at lipid species level during NETosis with class distinction and color-coded category assignments. Only lipid species which were significantly changing (p_adj_ < 0.05) in at least one of the comparisons are displayed. (f-n) Class-wise lipid regulation upon ionomycin treatment with examples of molecular lipid species for each class. (f-g) Upper panel displays fewer complex lipids such as fatty acyls, oxylipins and lysoglycerophospholipids. (i-k) Middle panel shows membrane lipids, major phospholipid classes (such as phosphatidic acid, phosphatidylcholines and phosphatidylethanolamines). (l-n) The last panel shows diacylglycerols, long-chain bases and phosphatidylinositols as potential signaling lipids. Examples have been provided for each panel. Fold changes for individual lipid species are shown as grey lines, the class average is depicted in color. The timepoints were always compared to the unstimulated quantities (Welch t-test, p_adj_ < 0.05 (n = 5 biological replicates).

The most significant change in complex phospholipids in PMNs is the upregulation of phosphatic acid (PA) peaking at 30 min (Fig. 2d), likely connected to phospholipase D (PLD) activity. As indicated by the decreasing levels of specific species, PC more than PE (Fig. 2 j, k) could be used for PA generation. It appears that PA regulation coincides with a diacylglycerol (DG) increase (at 2 hours), which is induced by diacylglycerol kinase (DGK) and phospholipase β/γ (PLCβ/γ). The DG increase is essential for neutrophil chemotaxis (*39*) and a key element in the production of complex GP and GL, an effect detectable across almost all species (Fig. 2l). Exclusive of the delayed esterase activity, oxidation of PUFAs via the 5/15-Lox or Cox-1/2 pathways becomes evident early on, leading to increased levels of HETEs and PGE2 (Fig. 2 e, g). Similar trends can be observed overall upon NET-induction when comparing the cellular to the released lipidome (Fig. 2 f-n), however, sphingolipid levels are substantially amplified indicating ER derived vesicular release (Fig. 2c). Whilst major sphingolipid classes (such as sphingomyelin and ceramides) remain unchanged (Fig. 2d left), sphingosine is increased within the cell, and S1P (LCBP) levels are elevated in the PMN release indicating sphingolipid signaling (Fig. 2 d, m) upon PMN activation via ionomycin, which is likely used to attract more neutrophils to the site of activation, fostering their adhesion and further degranulation (Fig. 2c, m) (*40, 41*). The magnitude of change is more prominent in the PMN release (Fig. 2b) with increased percentages (in mol) and numbers of regulated species (Fig. 2c), reflecting the intracellular developments. These findings highlight the complex and dynamic nature of neutrophil lipid metabolism during NET induction via ionomycin, and demonstrate their link to cellular function and inflammatory responses, underscoring the significance and potential impact of this research.

### Characterization of the lipidome upon NET induction with PMA

Compared to ionomycin, PMA stimulation (Fig. 3a) triggers less regulation (Fig. 3b, c, d), but also results in lower and delayed PLA_2_ activity in PMNs and the PMN release (Fig. 3d). Remarkably, almost no vesicular release can be observed after PMA stimulation. In PMNs, multiple lysolipid classes are elevated after 120 minutes with a simultaneous increase of fatty acyls (Fig. 3d, e). At the category level, only minor changes can be observed in PMNs (Fig. 3b) and major lipid classes like PC and PE remain overall unchanged. Lipid classes involved in glycerolipid synthesis, such as PG, PA, and DG, increase after 30 minutes (Fig. 3d, e, f, i, l), suggesting a similar core regulation in the lipidome as observed during ionomycin treatment. Interestingly, with PMA, the increased DG levels are utilized for the production of TGs and consequently, lipid droplet formation takes place between 60-120 minutes upon NET induction (Fig. 3d, e, n). The reduction of cholesteryl esters further indicates a comprehensive remodeling of storage lipids within PMNs (Fig. 3d, e, m). Compared to intracellular lipid levels, the lipid release upon PMA activation displays changes only within the first minutes, during which many lipid classes decrease. However, after 60 and 120 minutes, all released lipids are stabilized with no significant changes detectable at class level (Fig. 3c, d). Likely, the initial changes occur through stress-induced pinocytosis (*42*), commonly used for scavenging metabolites to meet any kind of energy demands. We also find fewer lipid shifts and one-fourth as many regulated lipids within the PMNs releasate compared with ionomycin stimulation (Fig. 3c). In addition, if we review NETosis from a structural lipid perspective, neither sn-nor double-bond positions are altered indicating no further stereochemical rearrangements during ionomycin or PMA treatment (Fig. S5).

**Figure 3.**
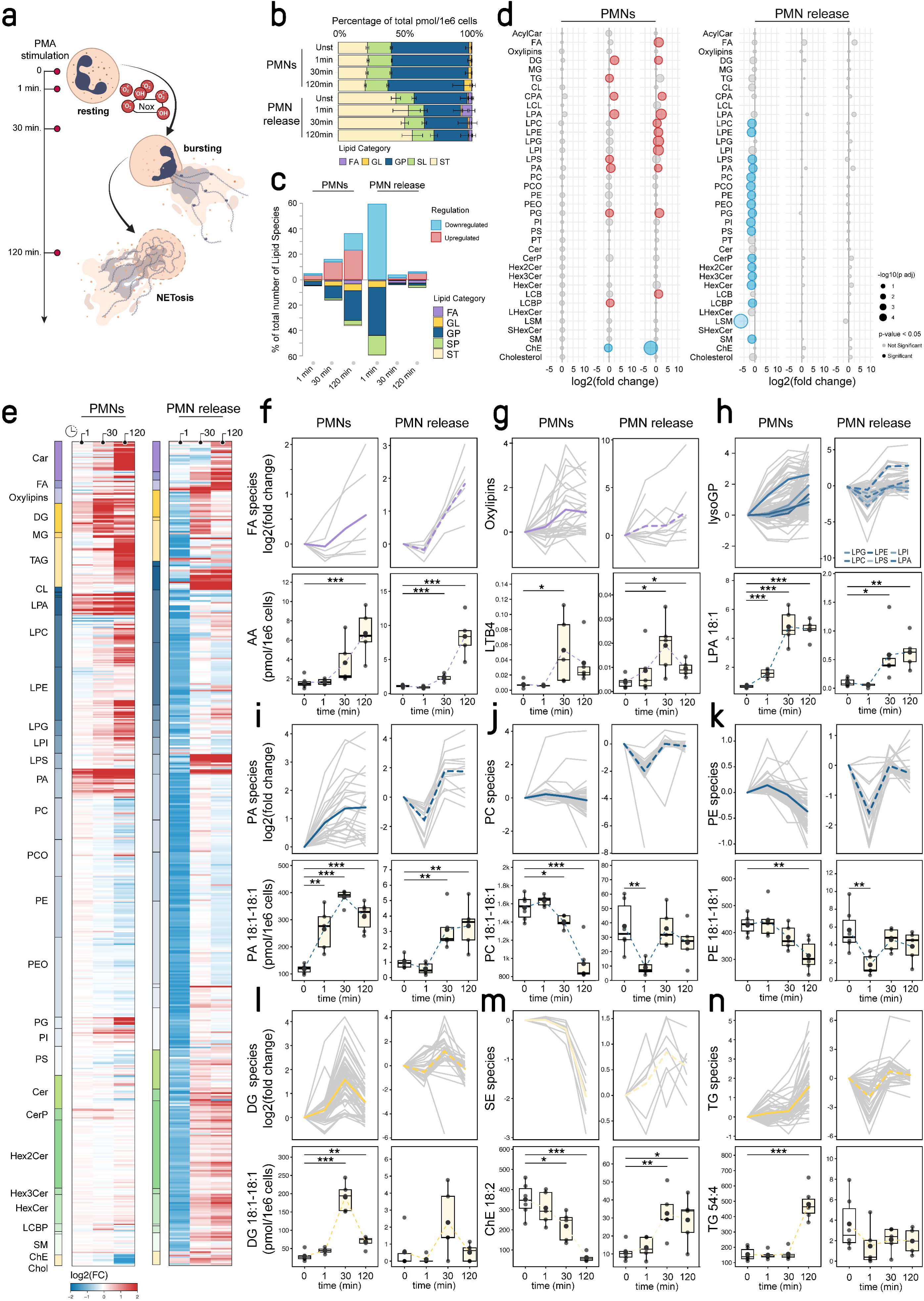
Dynamic gradual lipidome rearrangement of PMNs during NETosis triggered by diacylglycerols. (a) Timeline of sampling after NET induction using 100 nM phorbol ester (PMA). (b) Category-based lipidome remodeling of PMNs and release after NET-induction. (c) Quantitative assessment of the number and direction of species regulated by class during NETosis. (d) Ratio plots displaying class dynamics at 1, 20 and 120 minutes (left to right) for the cellular and the released lipidome. The ratios are displayed as log2 fold changes and depict considerable modulations of the PMNs and released lipidome at class level during NET-formation. Significant shifts are shown in color; blue for downregulated and red for upregulated classes. (e) Heatmap at lipid species level during NETosis at 1, 30, and 120 minutes. Only lipid species which were significantly changing (p_adj_ < 0.05) in at least one of the comparisons are displayed. Upper panel (f-h) displays less complex lipids such as fatty acyls, oxylipins and lysoglycerophospholipids. Middle panel (i-k) displays membrane lipids, major phospholipid classes such as phosphatidic acid, phosphatidylcholines and phosphatidylethanolamines, and the lower panel (l-n) shows diacylglycerols, sterols esters and triacylglycerols with the last two representing lipid droplet components. Examples are given for each of the panels. Fold changes for individual lipid species present as gray lines and the average per class is depicted in color. The timepoints were always compared to the unstimulated quantities (Welch t-test, p_adj_ < 0.05 (n = 5 biological replicates).

### NET induction is characterized by a time-dependent interplay of lipases

The lipidome shift during PMN activation is characterized by a combination of PLA_2_, PLC, and PLD activity (Fig. 4a), which, in the presence of ionomycin and PMA, triggers oxidative processes within the 5/15-lipoxygenase (Lox) or cyclooxygenase (Cox-1/2) pathways as indicated by the increased levels of oxylipins such as LTB4 or PGE2 (Fig. 2d, g and 3d, g). The activity of these lipases ultimately leads to elevated levels of phosphatidic acid (PA) and diacylglycerol (DG) (Fig. 4b), resulting in positive reinforcement through the protein kinase C (PKC) pathway.

**Figure 4.**
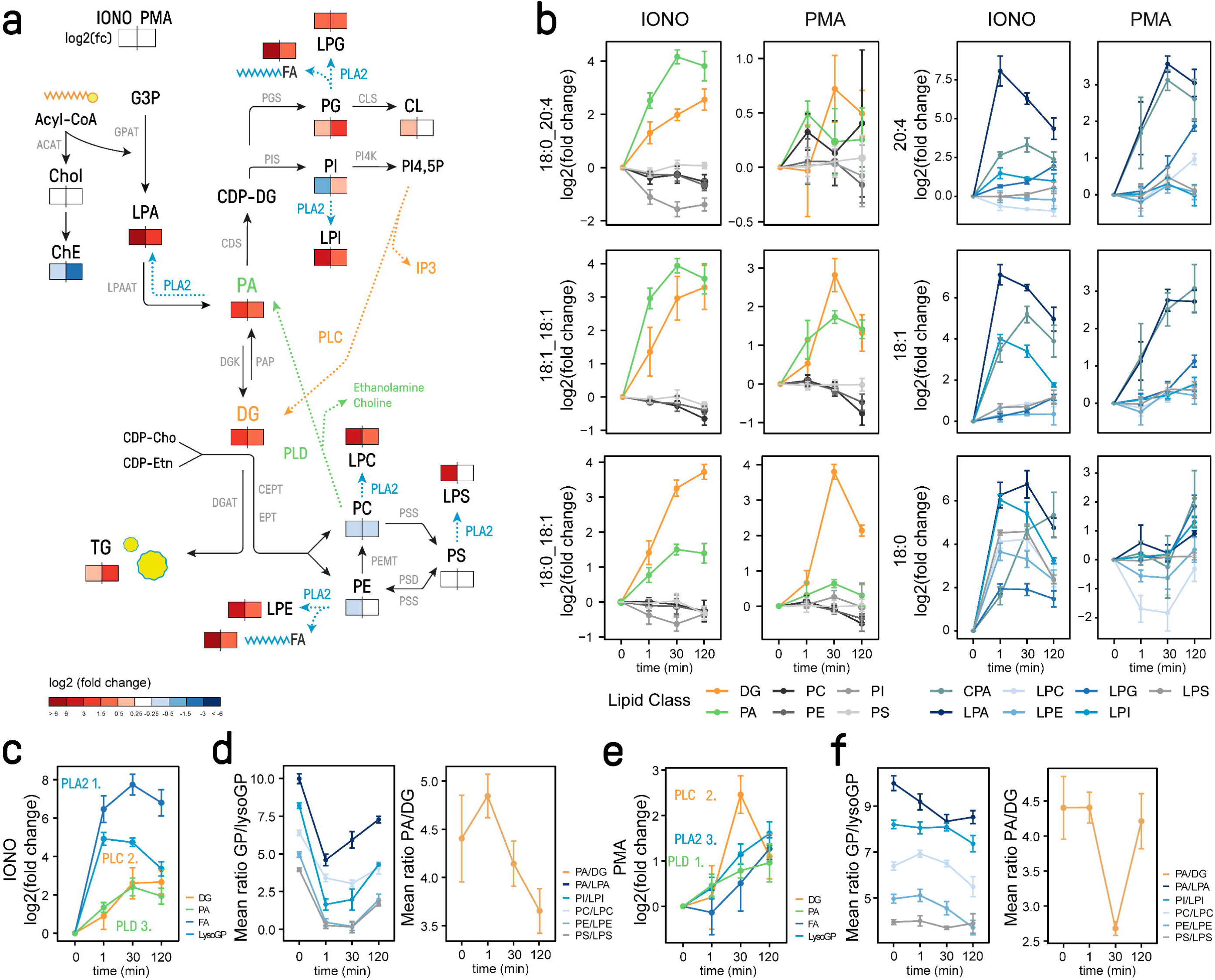
PMNs lipidome remodeling is characterized by interconnected phospholipase activity. (a) Monitored lipid metabolism and lipase activity during PMN activation using ionomycin (left) and PMA (right); significant changes are color-coded by log2 mean fold change scale compared to the control at 0 minutes; red indicates upregulated and blue downregulated lipid classes. (b) Interconnected lipid species during NETosis. Different species of diacylglycerols (DG, orange), phosphatidic acid (PA, green) and lyso-phospholipids (blue) are selected and compared to other phospholipid classes (grey). For PA and DG mono-, di- and polyunsaturated fatty acids and for the lyso-phospholipids saturated, mono-unsaturated and polyunsaturated fatty acids are displayed. (c) Summary of lipid class profiles reflecting PLA_2_ (blue), PLC (yellow) and PLD (green) activity during ionomycin treatment; class-level summaries of lyso-GP, FA (PLA_2_), DG (PLC) and PA (PLD) were used for readout. (d) The ratio of glycerophospholipids to lyso-glycerophospholipids is displayed indicating early PLA_2_ activity during ionomycin treatment, PA/LPA (green), PI/LPI (dark green), PC/LPC (violet), PE/LPE (grey), PS/LPS (pink); PA to DG ratio is reflecting onset of PLC versus PLD activity during PMN activation. (e) Summary of lipid class profiles reflecting PLA_2_, PLC and PLD activity during PMA treatment; class level summary of lyso-GP, FA (PLA), DG (PLC) and PA (PLD) were used for readout. PA to DG ratio reflects shifts in phospholipase activity between PLD and PLC during PMN activation (n = 5 biological replicates).

When enzyme activity is plotted using the relevant precursor-to-product ratios, the transient nature of lipase activity becomes evident (Fig 4 d, f). During ionomycin activation PLA_2_ activity rises quickly and reaches peak activity after 1 minute of stimulation (Fig. 4b), which is indicated by the steep increase of lyso-glycerophospholipids and fatty acids (Fig. 4b, c). PLD and PLC activity increases between 30-120 minutes, demonstrated by increasing levels of phosphatidic acid and diacylglycerol (Fig. 4b, c). As PLC levels continue to rise, PLD gradually declines towards the 120-minute mark.

In contrast, during PMA activation, PLD increases earliest, followed by PLC, and then PLA (Fig. 4f, e), which reaches its maximum activity at 120 minutes, while PLC activity decreases (Fig.4 b, e). Overall, both stimuli demonstrate at 120 minutes an overall elevated PLA_2_, PLD, and PLC activity, which is likely essential for the proper induction of NETosis.

### Effect of phospholipase signaling on in vitro NETosis and lipid dynamics

Subsequently, we addressed the physiological function of phospholipase signaling on NET formation, of which phospholipase D specifically has been identified as a crucial element in lipid modifications. We investigated the effect of pharmacological phospholipase D inhibition via specific inhibitor 5-Fluoro-2-indolyl deschlorohalopemide (FIPI) on NETosis in circulating human neutrophils by employing in vitro assays. The pretreatment of PMNs with FIPI evoked noticeable changes in cellular hallmarks of NET formation such as nuclear lobulation and plasma membrane rupture. In the absence of FIPI a clear increase in NET structures with a simultaneous decrease in nuclear lobulation was observed after ionomycin or PMA stimulation (Fig. 5a). In a concentration-dependent manner, pretreatment with FIPI resulted in an increased number of lobulated neutrophils and markedly decreased NET structures in activated PMNs, when compared to the solvent control (DMSO) (Fig. 5a), which is altogether indicative of reduced NETosis. Correspondingly, pretreatment with 1 µM FIPI resulted in a significantly impaired ionomycin- and PMA-induced NETosis compared to the solvent control, whereas FIPI treatment had no effect on unstimulated naïve PMNs (Fig. 5b).

**Figure 5.**
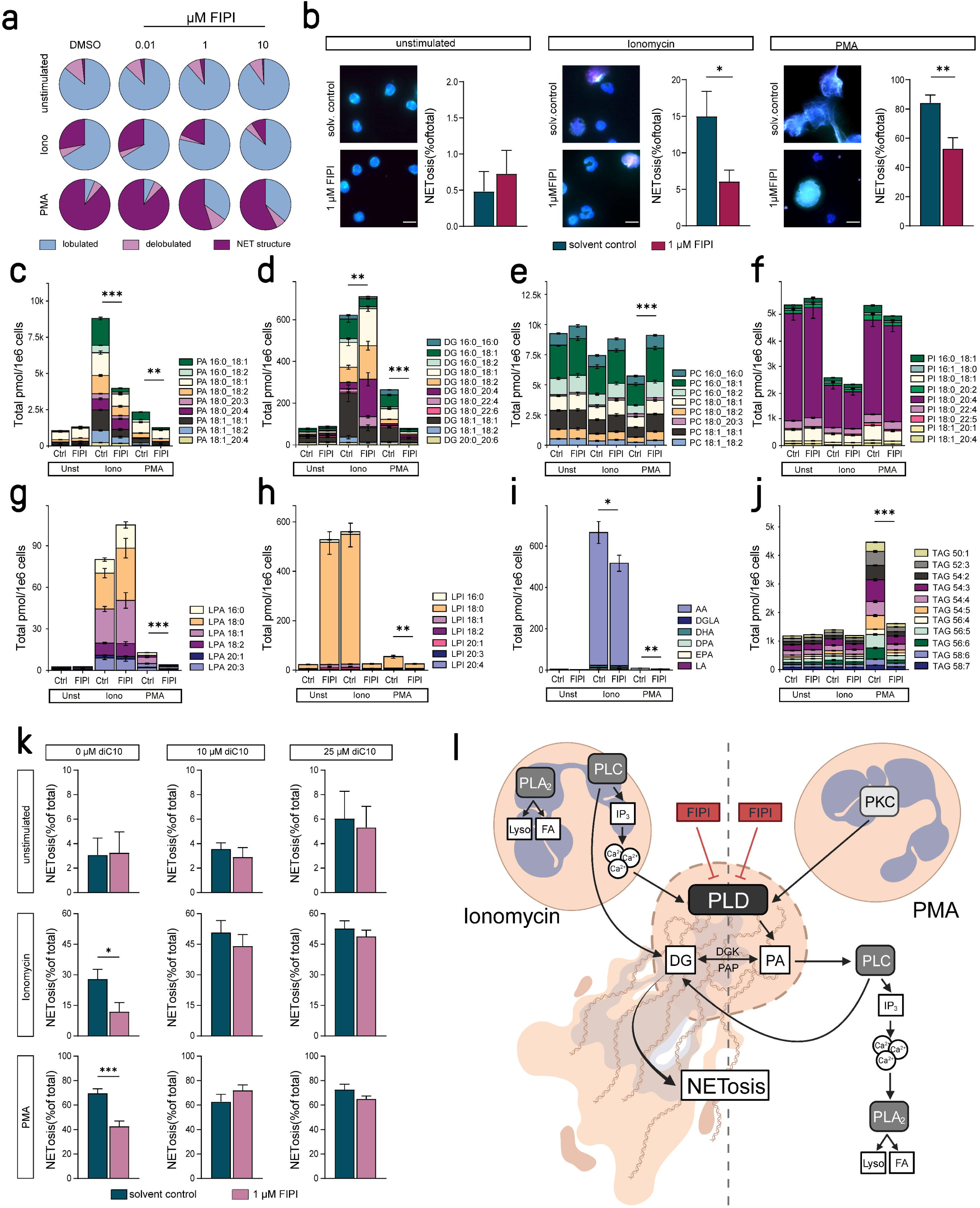
PLD inhibition reveals quantitative and qualitative traits of PA and DG signaling in NETosis. (a) Percentage of unstimulated, ionomycin (4 µM) or PMA (100 nM) stimulated PMNs at various cellular stages of NETosis (n=6) in the absence (DMSO) or presence of FIPI at the indicated concentration (b). Percentage of NETs (n=6) in unstimulated, ionomycin (4 µM) or PMA (100 nM) stimulated PMNs in the absence (solvent control) or presence of 1 µM FIPI. (b) Percentage of NETs (n=6) in unstimulated, ionomycin (4 µM) or PMA (100 nM) stimulated PMNs in the absence (solvent control) or presence of 1 µM FIPI. (c-j) dissected lipid class distribution visualized at species-level during PLD inhibition and NETosis triggered by PMA and Iono stimulation. The most abundant lipid species of the following lipid classes are depicted: (c) phosphatidic acid (PA), (d) diacylglycerols (DG), (e) phosphatidylcholines (PC), (f) phosphatidylinositol’s (PI), (g) lyso-phosphatidic acid (LPA), (h) lyso-phosphatidylinisitol (LPI), (i) fatty acids (FA) and (j) triacylglycerols (TG). The comparison of affected lipid classes is provided by corresponding controls. The data displays the average of n = 5 biological replicates, unpaired t-test p < 0.05. (k) Percentage of NETs (n=6) in unstimulated, ionomycin (4 µM) or PMA (100 nM) stimulated PMNs in the absence (solvent control) or presence of 1 µM FIPI and concurrent treatment with diC10 at the indicated concentrations (0, 10, 25 µM). (l) summary of the interconnected phospholipase network modulating the PMN lipidome and NETosis.

One of the identified hallmarks of lipid associated dynamics upon NET induction is the PLD-dependent generation of PA and DG (Fig. 4b). Thus, in a next set of experiments we investigated the effect of pharmacological PLD inhibition on the lipidome of PMNs. When compared to the solvent control, the presence of 1 µM FIPI lead to reduced PA levels upon ionomycin or PMA stimulation (Fig. 5c). Thereby, mainly mono-unsaturated and saturated PA species (e.g. PA 16:0_18:1) were affected by PLD inhibition (Fig. 5c) suggesting a corresponding substrate specificity. Notably, while in the presence of FIPI total DG levels were significantly downregulated after PMA stimulation, while upon ionomycin stimulation they remained unaffected (Fig. 5d). However, even with overall consistent DG levels, remodeling occurs in response to PLD inhibition after ionomycin treatment when compared to the solvent control (Fig 5d). Again, comparable to the affected PA species, mainly mono-unsaturated and saturated DG species emerging from mono-unsaturated and saturated PA species were downregulated upon FIPI treatment and ionomycin-induced NETosis (Fig. 5d). In addition, DG lipid species containing the same PUFA released by PLA_2_ are increased (Fig. 5d), which indicates DGK activity via PLC shunt compensating for the loss of DG containing saturated and mono-unsaturated such as DG 18:1_18:1, DG 16:0_16:0 and 16:0_18:1. Adding the PLD inhibitor validates our point that PA is PC-derived, as shown by the restoration of PC levels after FIPI treatment (Fig. 5e). Interestingly, PLA_2_ activity also seems to be influenced by PLD inhibition as reflected by the reduced LPA, LPI and FA levels during PMA-induced NET-activation (Fig. 5g-i), which points towards an interconnected downstream phospholipase network. In contrast, PLD inhibition during ionomycin induced NET activation affects less PLA_2_ activity, as evidenced by the fact that PI levels are not restored (Fig. 5f) – indicating that PLA_2_ function is maintained under these conditions. Additionally, an effect on triacylglycerol synthesis and consequently lipid droplet production was observed in the PMA-stimulated PMNs after PLD inhibition with TG levels not changing upon PLD inhibition (Fig. 5j).

PA and particularly mono-unsaturated and saturated DG species are well-established upstream effectors of protein kinase C (PKC), which plays a pivotal role in ionomycin-as well as PMA-induced NETosis (*43*). Moreover, since FIPI treatment in ionomycin- and PMA-induced NETosis resulted in markedly decreased mono-unsaturated and saturated DG species, we investigated the effect of diC_10_ treatment on in vitro NET formation in PLD-inhibited neutrophils. As depicted in figure 5k, the treatment with diC_10_ concurrent with PLD inhibition completely restored NETosis after ionomycin or PMA stimulation (Fig. 5k), demonstrating the importance of PA and DG-induced signaling in the process of NET formation (Fig. 5i), which ultimately underscores our suggested model of action (Fig. 5l).

## Discussion

NETs are linked not only to infectious diseases, but also affect the development and progression of thrombo-inflammatory pathologies such as thrombosis (*4, 44, 45*) or ischemic and inflammatory heart diseases (*6–8*) and are additionally connected to cancer (*10*) and diabetes (*9*), all of which contribute significantly to morbidity and mortality worldwide (*46*). Consequently, NETosis is on a pathophysiological level highly relevant and potential clinical applications for prophylaxis as well as treatment of NET-related disorders remain underdeveloped. Unfortunately, the exact cellular and molecular signaling mechanisms underlying NETosis are insufficiently defined. Particularly, the quantitative lipidome and the role of dynamic lipid alterations during NETosis is often neglected, although the lipidome of neutrophils seems to be distinct (*22*) and a broad range of neutrophil functions such as phagocytosis, adhesion, secretion of microbicidal compounds and NETosis are crucially dependent on the properties and lipid composition of the plasma membrane. Therefore, in the underlying study we employed state-of-the-art lipidomics workflows to characterize for the first time the quantitative lipidome of human neutrophils and its dynamic alteration during ionomycin- or PMA-induced NETosis. This distinct and innovative approach uncovered a lipidomic dynamic range spanning nine orders of magnitude in naïve neutrophils with ether-linked lipid species representing the predominant headgroup type (Fig. 1), which is in accordance with previous reports (*22*) and corroborates our methodological approach. Additionally, the lipidome comprises a highly dynamic range (nine orders) and an impressive variety of components (1,039 species, across 37 classes), indicating a large functional diversity among lipids. The remarkably large proportion of hexosyl and di-hexosylceramides and the relatively low proportion of cholesterol and SM indicate a rather flexible membrane. In addition, we characterized the dynamic changes in the lipid landscape during NETosis. Distinctive, stimulus-dependent patterns of membrane remodeling were observed after ionomycin and PMA activation, however, the main lipid classes modified including PA, DG, lyso-GP and fatty acyls proved similar. In this context, our study identified phospholipase lipase networks and PLD specificity towards saturated and monounsaturated fatty acyls as crucial elements of in vitro NETosis upon stimulation with ionomycin or PMA and altogether established a comprehensive and detailed reference lipidome for circulating naïve neutrophils.

PLD is strongly expressed in immune cells including neutrophils (*47*) and its activation is implicated in phagocytosis and oxidant production (*48*). PLD predominantly generates the anionic phospholipid PA, which is known to alter the lipid composition of the plasma membrane’s inner leaflet causing a negative membrane curvature. Interestingly, although ionomycin- and PMA-induced NETosis manifest as distinct lipidomes and lipid dynamics, both inducers converged on the generation of PA and DG (Fig. 4a). In other immune cells, modified membrane curvatures caused by PA are known to directly impact cellular shape and exocytotic and phagocytotic processes (*49*). Membrane curvature was demonstrated to serve as key regulator of the spatial distribution of the ubiquitously expressed mechanosensitive ion cannel Piezo1 in living cells (*50*), which was just recently identified as a positive regulator of shear-induced NETosis (*51*). Thus, at least under shear stress, PA-induced changes in plasma membrane curvature may directly contribute to the increased plasma membrane permeability and rupture observed at distinct stages of NETosis (*18*). Nevertheless, further studies are required to determine the exact role of PA and mechanotransduction during NETosis, considering the neuronal-specific Piezo2 channel seems to be negatively affected by endogenous PA (*52*). Intriguingly, in the past years the involvement of NETs in cancer and diabetes has been well-acknowlegded (*9, 10*), including reports on the major role of PLD in metastasis and chemoresistance (*53*) and metabolic disorders such as diabetes (*47, 54*), underscoring the importance a PLD-dependent signaling mechanism underlying NETosis.

Besides affecting physical plasma membrane properties, PA also serves as an important signaling lipid in the plasma membrane (*55*). and its protein binding properties are affected by cholesterol (*56*) a sterol known to affect NETosis (*20*) and which is the most abundant molecule in naïve neutrophils (Fig. 1c, bottom). The importance of PA protein binding properties is further corroborated by the observation that PA binds directly to actin-related protein 3 (Arp3), thus affecting actin cytoskeleton dynamics and subsequently leukocyte adhesion (*57*) whereas especially the early and quick rearrangement of the actin cytoskeleton represents a defined major hallmark of NETosis (*18*).

Both PA and DG reside in an equilibrium via PAP and DGK driving PKC activation. Moreover, remarkably PLD activation seems to affect the release of ROS from neutrophils upon PMA stimulation (*58*) a mechanism most likely involving protein kinase C (PKC)-dependent signaling (*59, 60*). Thereby, in an isoform-dependent manner, PKC signaling is undoubtedly linked to the formation of NETs upon calcium ionophore and PMA stimulation (*17, 43, 61*). The involvement of PLD-dependent signaling in the regulation of PKC activity and its localization in immune cells is increasingly evident, since the simultaneous generation of PA and DG is essential for the translocation of a specific PKC isoform to the plasma membrane (*62*). Our results demonstrated that the pharmacological inhibition of PLD by FIPI impaired ionomycin- and PMA-induced NETosis in vitro in a concentration-dependent manner (Fig 5a, b). More remarkably, this effect was completely reversed in the presence of the DG analog and PKC activator sn-1,2-didecanoylglycerol (diC_10_) (Fig 5k), which is in accordance with data showing a restored oxidase activation in neutrophils upon PLD inhibition (*48*). These findings most likely point to the involvement of generated PA and DG in PKC signaling during ionomycin- and PMA-induced NETosis. However, further investigations are required to unravel the exact role of PA- and DG-induced NETosis, since diC_10_ could be transformed to PA via a DGK-dependent mechanism.

Conclusively, this study delineates for the first time comprehensively the quantitative lipidome of naïve neutrophils and more importantly the dynamic lipid alterations underlying NETosis upon iono- and PMA stimulation. Furthermore, PLD-dependent PA and DG generation was clearly linked to NETosis. Thus, the presented results unravel new mechanistic insights into the molecular signaling of NETosis, which will contribute to our understanding of the multifaceted nature of NET formation and its connection to clinically relevant human disorders. From a translational approach, our results demonstrate that pharmacological PLD inhibition or lipid-associated manipulation of NETosis may be a promising new strategy for the prophylaxis and treatment of NET-associated inflammatory diseases.

## Supporting information

Supplemental Results

## Acknowledgments

The authors thank Daniela Eißler for excellent technical assistance and acknowledge the support by the Open Access Publishing Fund of University of Tübingen.

## Authorship

O.B. and R.A. developed the concept. P.M., C.C., O.B., and R.A. designed, conducted, and analyzed the experiments. P.M., C.C., G.D.L., N.N.T., J.A.M., R.S.E.Y., S.R, J.M., J.A., M.F., N.S., F.K., M.Z., C.H. and S.R.E. conducted experiments. P.M., C.C., S.E. and R.A. analyzed data. C.C. reviewed the methodological aspect with P.M., R.A. and O.B. All authors wrote and revised the manuscript.

