## Supplemental Results for "Dynamic neutrophil lipidome remodeling during induction of NETosis"

### Supplementary Results

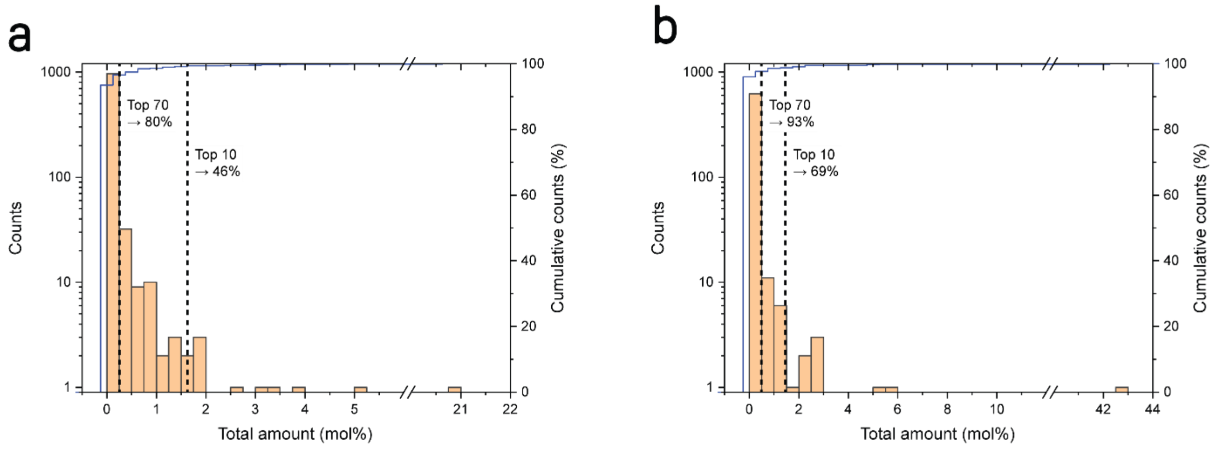


**Figure S1: The PMN resting lipidome is complex and diverse**. (a, b) Cumulative analysis of lipid abundance. The lipids are plotted according to their abundance, the Top 10 and Top 70 intervals are displayed. The PMN lipidome (a) is compared against the less complex release of PMNs (b). The left *y* axis displays the number of lipids, the right one the cumulative counts the total amount in mol is given at the *x* axis.

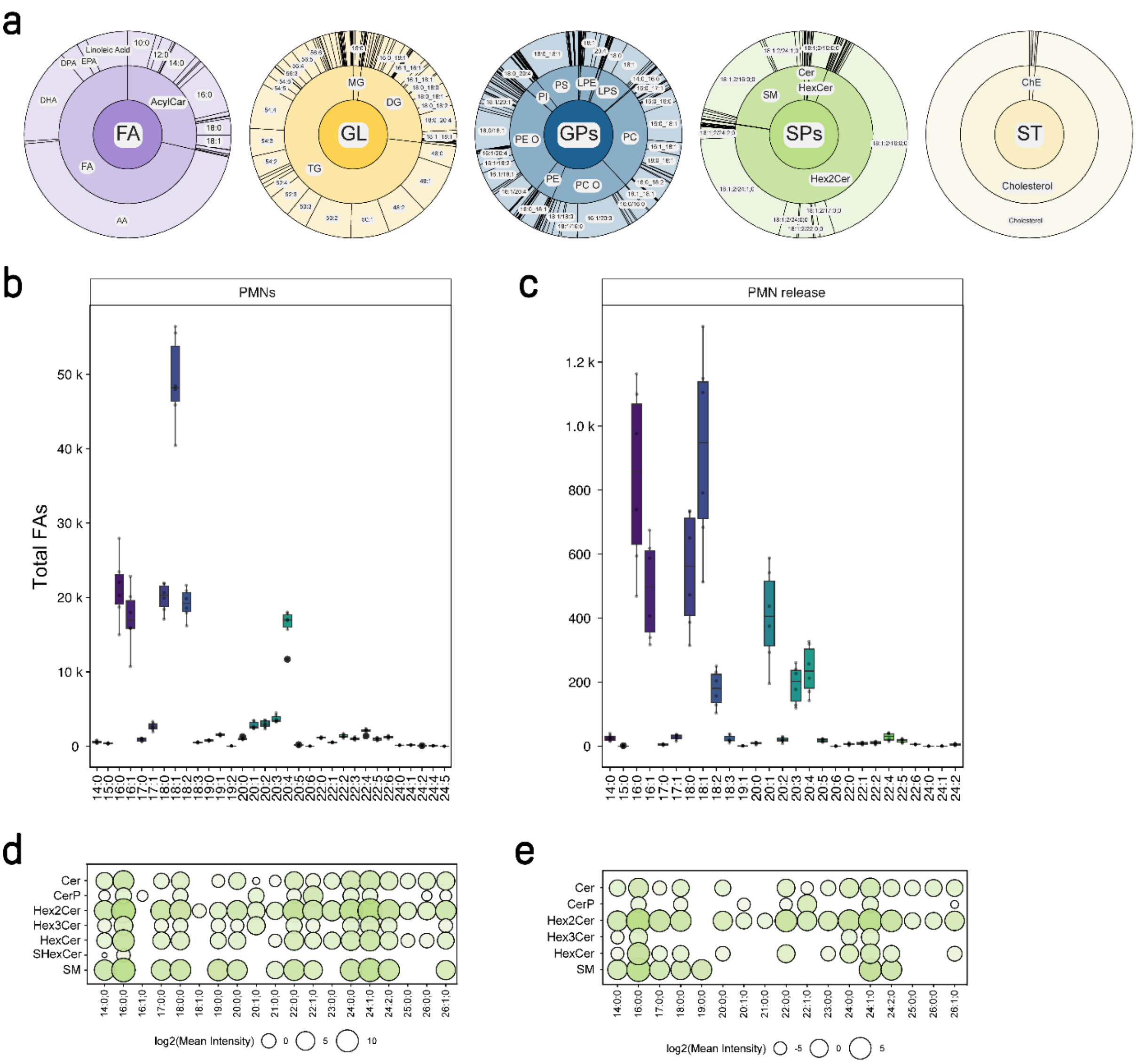


**Figure S2: Fatty acyl distribution based on MS2 experiments of the release and PMN pellet.** (a) Displays the category and class based fatty acyl distribution of the PMN release. (b,c) Displays the fatty acyl distribution across the lipidome, thereby release (c) and cell can be compared (b). (d,e) Fatty acyl distribution of sphingolipids in PMNs and the release without any stimulation. All measurements are based on 5 biological replicates.

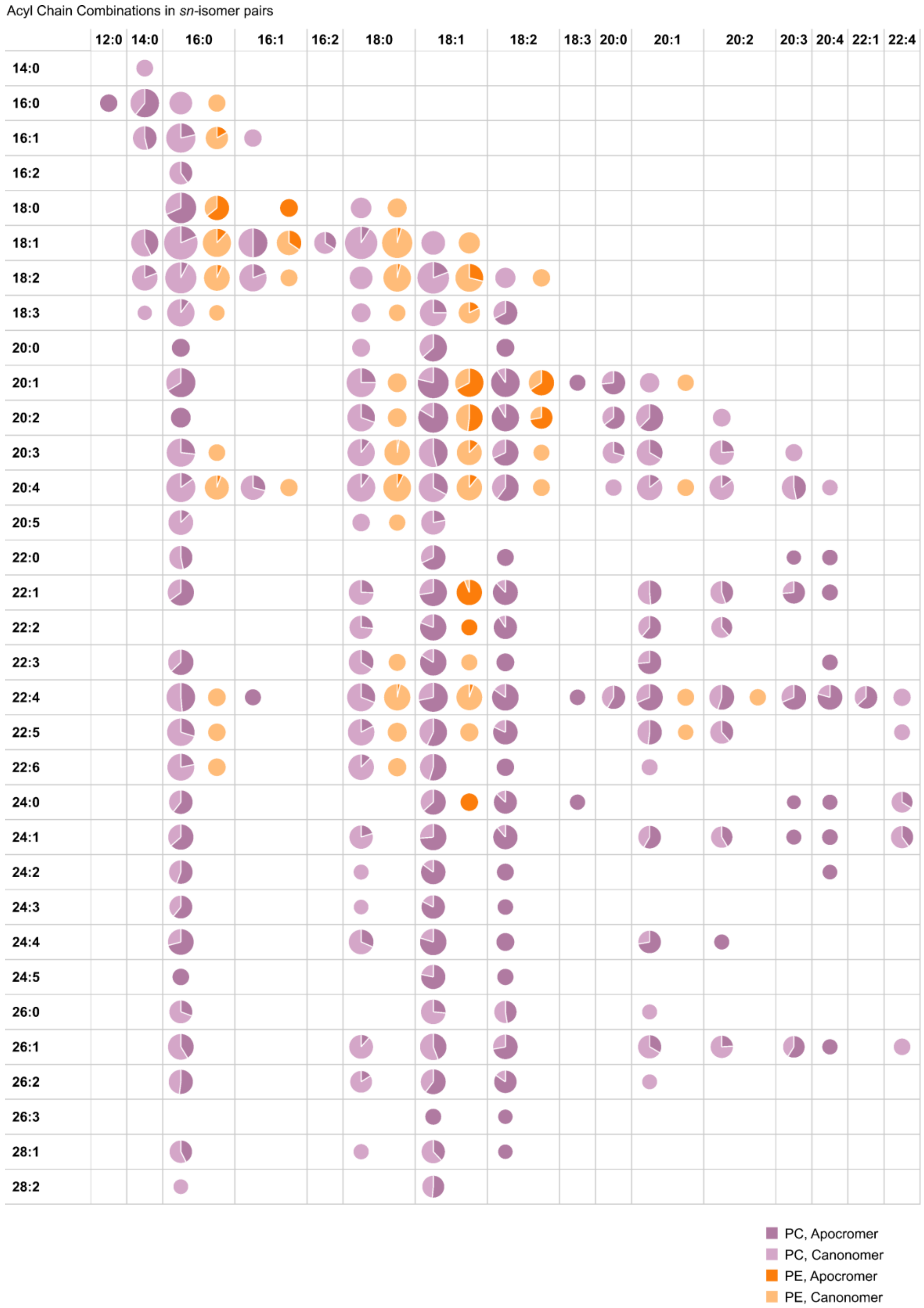


**Figure S3: sn positional analysis of PCs and PEs of human unstimulated PMNs**. Analysis of PC and PE molecular lipid species with *sn*-position structural detail detected at the MS3 level with CID/OzID, separated by component acyl chains. Each pie slice reflects sum intensity of relevent MS3 fragments for three replicates of PMNs (unstimulated). Chart is limited to even-numbered carbon chain lengths.

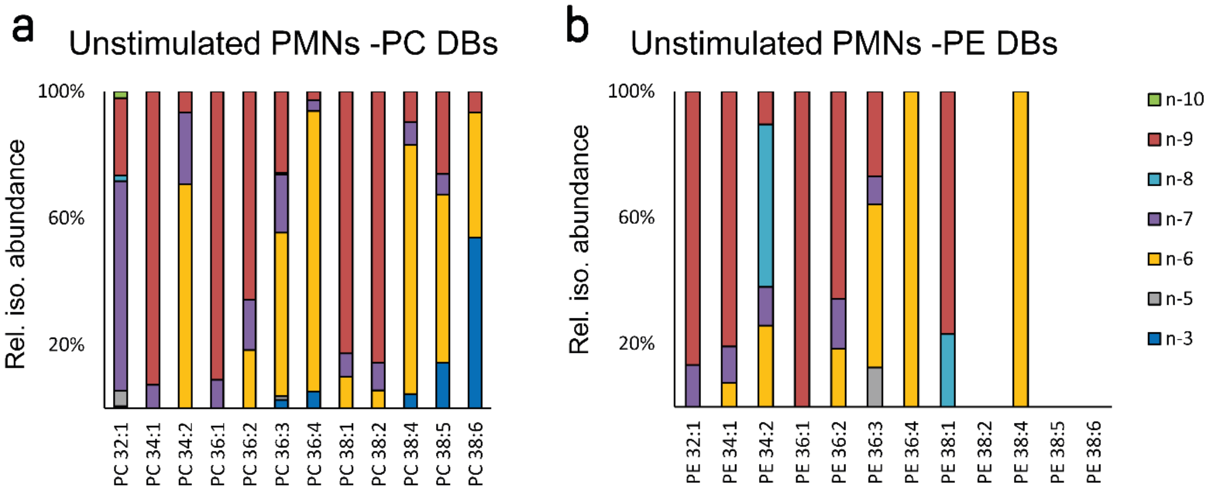


**Figure S4: Double bond isomer analysis of 20 abundant PC and PE lipids from unstimulated PMNs.** Data were acquired using ozone-induced dissociation (OzID) and as such require a correction coefficient that accounts for the variation in ozonolysis rate kinetics between carbon-carbon double bond positions to be truly reflective of mol% abundance. Double bond isomers shown as n-3 (dark blue) and n-6 (yellow) are PUFAs with methylene interrupted double bonds (i.e., n-6,9,12). All remaining DB positions describe various MUFA isomers. (Mean average displayed, n=3).

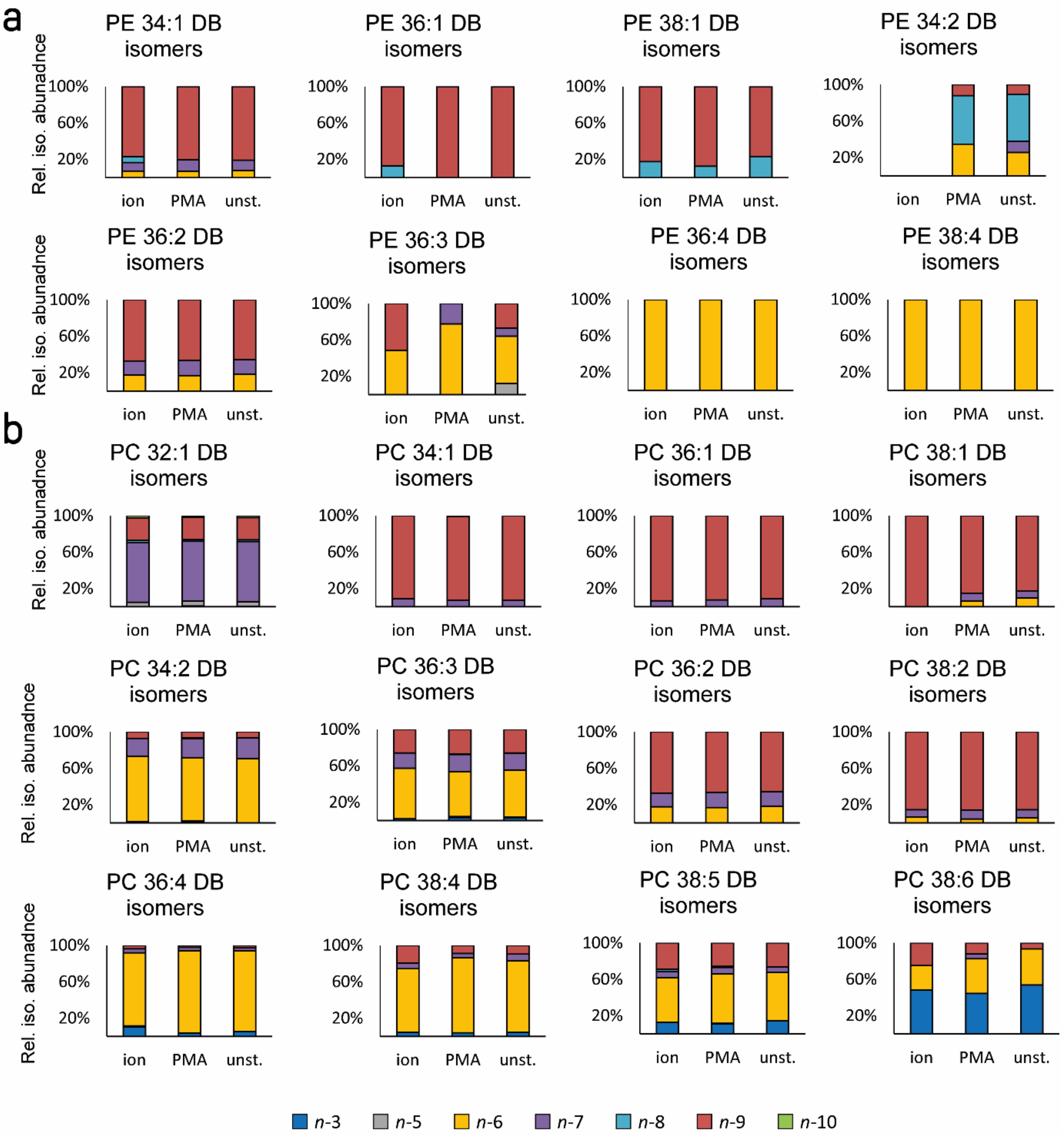


**Figure S5: Double bond isomer analysis of 20 abundant PEs and PCs from stimulated and unstimulated PMNs.** Data were acquired using ozone-induced dissociation (OzID) and as such require a correction coefficient that accounts for the variation in ozonolysis rate kinetics between carbon-carbon double bond positions to be truly reflective of mol% abundance. Double bond isomers shown as n-3 (dark blue) and n-6 (yellow) are PUFAs with methylene interrupted double bonds (i.e., n-6,9,12). All remaining DB positions describe various MUFA PC isomers. (Mean average displayed, n=3).
